# Slc7a11 modulated by POU2F1 is involved in pigmentation in rabbit

**DOI:** 10.1101/607978

**Authors:** Yang Chen, Shuaishuai Hu, Lin Mu, Bohao Zhao, Manman Wang, Naisu Yang, Guolian Bao, Cigen Zhu, Xinsheng Wu

**Affiliations:** College of Animal Science and Technology, Yangzhou University, Yangzhou, Jiangsu, China; Joint International Research Laboratory of Agriculture & Agri-Product Safety, Yangzhou University; Animal Husbandry and Veterinary Research Institute, Zhejiang Academy of Agricultural Sciences, Hangzhou, Zhejiang, China; Jinling Rabbit Farm, Nanjing, Jiangsu, China

**Author notes:** Corresponding Author Xinsheng Wu, College of Animal Science and Technology, Yangzhou University, 48 South University Ave, Yangzhou, Jiangsu, 225009, P. R. China.

**Keywords:** Rex Rabbit, melanocyte, pigmentation, Slc7a11, POU2F1

## Abstract

Solute carrier family 7 member 11 (Slc7a11) codes for a cystine/glutamate xCT transporter and can control production of pheomelanin pigment to change fur and skin colors of animals. Previous studies found that the skin expression levels of Slc7a11 varied significantly with the fur colors of the Rex Rabbit. However, it is not yet known the molecular regulation mechanism of Slc7a11 in pigmentation. Here rabbit melanocytes were first isolated and identified. The distribution and expression pattern of Slc7a11 was confirmed in rabbit skin with different fur colors. Slc7a11 could affect the expression of pigmentation related genes and thus affect melanogenesis. Meanwhile, Slc7a11 decreased melanocytes apoptosis, but inhibition of Slc7a11 enhanced apoptosis. Furthermore, it was found that POU2F1 protein bound to the -713 to -703 bp region of Slc7a11 promoter to inhibite its activity by dual-luciferase reporter and site-directed mutagenesis assay. This study uncover the function of the Slc7a11 in melanogenesis and provided in-depth analysis of the mechanism of fur pigmentation.

## Introduction

The fur color of mammals mainly depends on melanin deposition, and melanogenesis is mainly regulated by melanocytes. The production of different types of melanin by melanocytes, together with different a distribution of these pigments, result in a variety of hair colors in mammals (Slominski et al., 2004). Related genes, such as TYR, TYRP1, ASIP, MITF, and CREB1, have been found to regulate melanin deposition (Hartman and Czyz, 2015; Hida et al., 2009). Previously, by using the transcriptome sequencing (RNA-Seq), a significant difference was found in the expression of the Slc7a11 gene in the skin of rabbits with different fur colors. It was speculated to be involved in fur pigmentation (Qin et al., 2016).

In the melanogenesis pathway, both eumelanin and pheomelanin are derived from a common precursor named dopaquinone (Ito, 2006). Cystine or glutathione is required for the production of pheomelanin, and it is xCT, the protein encoded by the Slc7a11 gene, that acts as a vector to transport extracellular cystine into the cell and maintain normal intracellular glutathione levels. Pheomelanin and eumelanin together form a mixed pigment that determines the skin and fur color of animals (Kim et al., 2001; Sato et al., 1999; Shih and Murphy, 2001). In the hair of Slc7a11 gene-mutated mice (sut), the level of pheomelanin was significantly decreased, while the eumelanin level was substantially unchanged, so that the wild-type mice with yellow background appeared gray (Chintala et al., 2005). The sut mutation results in a huge deletion in the Slc7a11 gene, but similar deletions could not be found in this region of Rex Rabbits with six different fur colors, namely black (BL), chinchilla (CH), white (WH), brown (BR), protein yellow (PY), and protein chinchilla (PC). SNPs in the exon region of Slc7a11 were also scanned, but no mutation site was found. This indicates that Slc7a11 is highly conserved in the population (data not shown). Currently, studies on the functions of Slc7a11 mainly focus on its important roles in cell proliferation (Liu et al., 2007), oxidative stress response (Bridges et al., 2001), and Alzheimer’s disease target treatment (Qin et al., 2006). Research studying its regulatory mechanisms is focused on microRNAs affecting cancer development and apoptosis by targeting and regulating Slc7a11 (Liu et al., 2007; Wu et al., 2017). Few studies regarding melanin deposition have been reported.

To explore the molecular regulation mechanism of Slc7a11 in the melanin deposition of Rex Rabbit fur, rabbit melanocytes were isolated and identified. The expression pattern of Slc7a11 in Rex Rabbit with different fur colors was analyzed. Further, it’s verified that POU2F1 had an important regulatory role in the transcriptional activation of the Slc7a11 gene promoter. This result provides a theoretical basis for further analysis of the deposition mechanism of the fur pigmentation as well as for the transformation of fur color in animals.

## Materials and Methods

### Primary separation and culture of rabbit melanocytes

Rabbit was injected with anesthetic on the back, and a piece of the back skin (1.5 cm x 1.5 cm) was dissected. Any particles on the skin surface and subcutaneous connective tissue were removed. The skin sample was digested with 0.25% DispaseII enzyme digestion solution (Sigma) for 14-16 hours at 4 °C. The epidermis was gently peeled off from the dermis, cut into small pieces, and digested with 0.25% trypsin (Gibco) for 8 minutes at 37 °C. The sample was then filtered through a 200 mesh filter and the supernatant discarded. The cells were resuspended in M254 medium (Gibco) and incubated at 37°C in a 5% CO2 incubator. The cells were digested with 1 mL of 0.25% trypsin and subcultured.

### DOPA staining

Melanocytes in the logarithmic growth phase were used to prepare sterile cell culture slides. The inoculated 24-well plates were cultured for 3 days and treated with 1 mL of 4% paraformaldehyde fixative (Solarbio) for 30 minutes at 4°C. The plates were washed 3 times with pre-cooled PBS prior to the addition of L-DOPA (Sigma). After incubation in L-DOPA for 1 day, the incubation solution was renewed and the plates were further incubated at 37 °C for 12 hours, with constant observation once every 30 min. The plates were washed with PBS once the staining was complete and observed under a microscope.

### Immunostaining

Melanocytes in the logarithmic growth phase or skin tissues of Rex Rabbits with 6 different fur colors were used to prepare slides. The slides were incubated with primary antibodies Slc7a11 (1:500 rabbit polyclonal, Abcam), S-100 (1:500 mouse monoclonal, Boster), TYRP1 (1:250 rabbit polyclonal, Abcam), TYR (1:1000 rabbit polyclonal, Abcam) overnight at 4°C, with PBS as a negative control. The slides were subsequently incubated with IgG secondary antibody (1:2500 goat polyclonal, Abcam) at 37°C for 20 minutes and developed for 3∼5 minutes at room temperature in the dark with freshly prepared DAB solution (Boster). The slides were observed under a microscope.

### RACE and cloning of Slc7a11 gene

Three specific 5’ RACE primers and two 3’ RACE primers were designed according to the Race kit instructions (Invitrogen & Clontech) (Table S1). The full-length cDNA sequence of the Slc7a11 gene was assembled based on known sequences and 5’ and 3’ RACE results, and submitted to NCBI (Accession no.: KY971639.1). The Slc7a11 cDNA was reconstructed into the pEGFP-N1 vector with restriction enzymes *Hind*III and *Sac*II.

### Knockdown of Slc7a11 by siRNA

Fluorescently labeled siRNAs (with 5’ FAM modification) and Negative Control siRNAs were purchased from Shanghai GenePharma Co., Ltd (Table S2). When the melanocytes confluence reached about 65%, the siRNA oligo / Lipofectamine™ 2000 (Invitrogen) complex at a ratio of 1:2 was prepared for transfection. After 24 hours, the transfection efficiency was examined by fluorescence microscopy.

### Real-time PCR

Real-time PCR was carried out using GAPDH as an internal reference. The fluorescent quantitative primers for the detection genes are shown in Table S3, and each sample was repeated 3 times. The relative expression of the target gene was calculated by the _ΔΔ_Ct method; namely, the fold difference between the target gene and the reference gene (experimental group)/the fold difference between the target gene and the reference gene 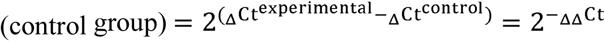

### Simple Western analysis

The pre-cooled RIPA lysis buffer (Sigma) was mixed with PMSF (with a final concentration of 1 mM) and added to the tissue or cell samples, which were centrifuged at 10000 rpm for 5 minutes at 4°C. The supernatant was discarded and the total protein obtained. Simple Western analysis was performed using the Wes Simple Western (Protein Simple) system. The test results were analyzed using the Compass program.

### Apoptosis assay

The cell apoptosis rate was measured with the Annexin V-FITC Apoptosis Detection Kit (Vazyme, China), according to the manufacturer’s instructions. Cells were sorted by fluorescence-activated cell sorting using the Flow cytometer FACSAria SORP (Becton Dickinson, USA).

### Determination of melanin level

The cells were lysed with 1 mL 0.2 mol/L NaOH. The cell lysate was collected and incubated at 37 °C for 2 hours. Wavelength measurement was performed at 475 nm using a microplate reader. The standard curve was plotted using the Melanin synthetic standard (Sigma). Each group was repeated 3 times, from which the melanin level was calculated.

### Luciferase vector construction and reporter assays

Promoter-specific primers were designed using O1igo7 (Table S4), and the Slc7a11 promoter region was analyzed using PROMO (http://www.cbs.dtu.dk/services/Promoter/) to obtain possible transcription factor binding sites. The Slc7a11 promoter was reconstructed into the pGL3-basic vector with restriction enzymes KpnI and BgIII. The internal reference plasmid pRL-TK and the recombinant plasmid were co-transfected into RAB-9 cells (ATCC), with the pGL3-basic plasmid and pRL-TK plasmid co-transfected cells as the negative control group and the cells with no substance transfected as the blank group. The transfected cells were collected and analyzed using the Dual-Luciferase Reporter Assay System (Promega).

### Electrophoresis mobility shift assay (EMSA)

The nuclear proteins of melanocyte were extracted and the concentrations determined. Based on the binding sequence of POU2F1, normal and mutant probes were designed (Table S5) and biotinylated at the 5’ end. The EMSA reaction system was formulated as shown in Table S6. The cold-competitive EMSA reaction system is shown in Table S7. The samples were analyzed by Native-PAGE, transferred, and UV cross-linked prior to carrying out the chemiluminescence reaction, development, and photographing.

### Statistical analysis

Each experiment was repeated at least three times and statistical significance between experimental and control groups was analyzed by Independent-Sample Test and one-way ANOVA. The results are presented as mean ± standard deviation (SD) at two levels of significance, *P* < 0.05 and *P* < 0.01.

## Results

### Separation and identification of rabbit melanocytes

The back skin of black Rex Rabbits was collected and cells separated by a two-step enzyme digestion method. After 12 hours of isolation, the keratinocytes were observed with a cobblestone-like appearance and accounted for the majority of cells. The melanocytes, which had the unique bi-polar dendritic morphology, were small in number. However, as cells grew, the keratinocytes gradually died out, the melanocytes continued to divide, and the cell culture became purer. When the cells were passed to the third generation, the keratinocytes were almost absent. The melanocytes were dominant with special growing follicles and strong refraction (Figure 1a). The melanocyte marker genes MITF, TYR, and TYRP1 were detected by semi-quantitative PCR (Figure 1b). The isolated cells stained with L-DOPA staining contained brown or black particles (Figure 1c). Immunocytochemical staining of S-100, TYR and TYRP1 revealed that the markers were expressed in the melanocytes. Compared with the negative control, S-100 staining showed the cytoplasm and dendrites were positively stained brown. The nucleus was brownish yellow in the TYR staining, and light brown in the TYRP1 staining (Figure 1d). This indicated that the rabbit melanocytes were successfully isolated and identified, providing experimental materials for this study.

**Fig. 1.**
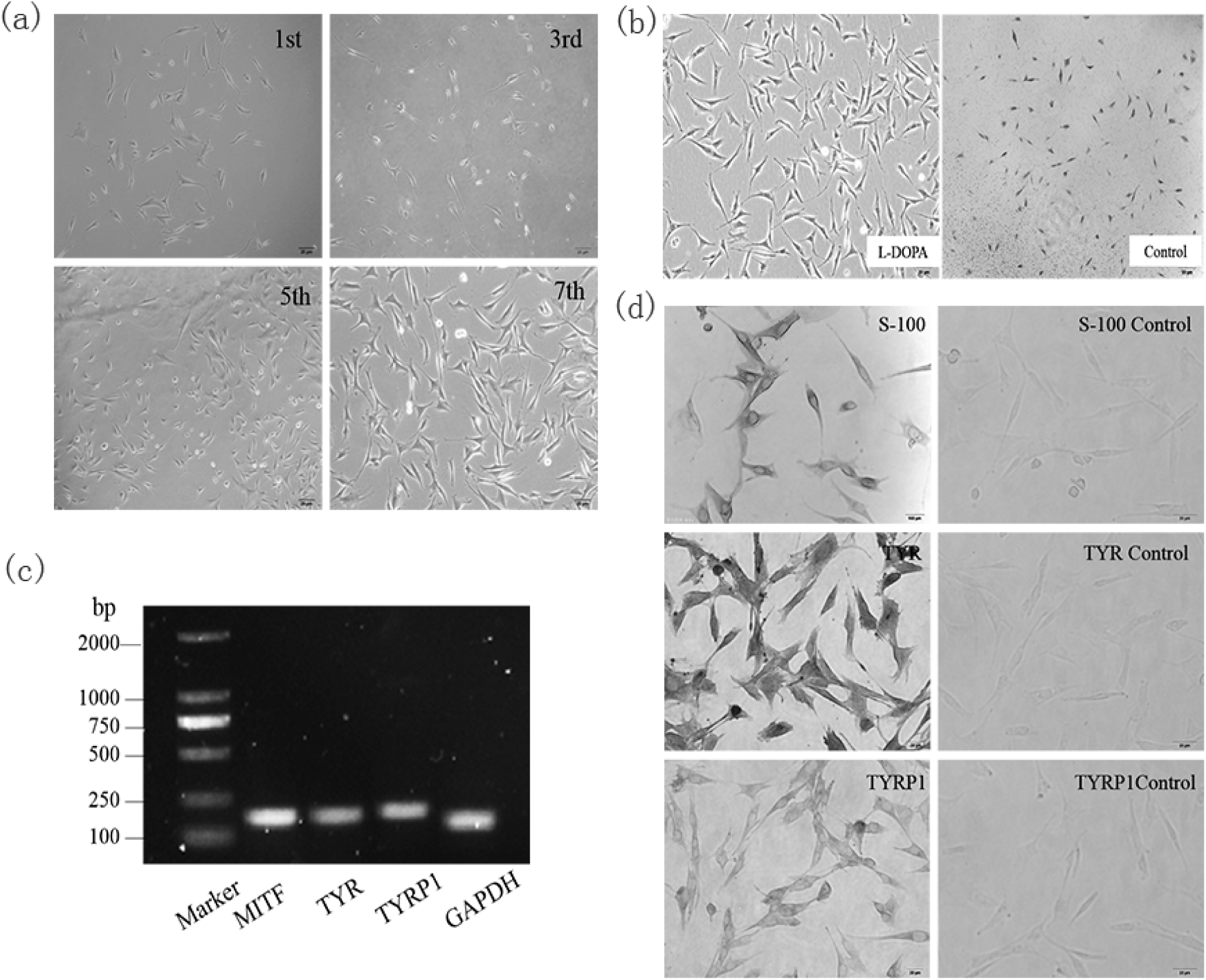
The separation and identification of melanocytes of rabbits. (a) Morphology of the 1^st^, 3^rd^, 5^th^, and 7^th^ generation melanocytes isolated by the two-step enzyme digestion method (100×). After 12 hours of isolation and culture, cobblestone-like keratinocytes and bi-polar dendritic melanocytes were observed. As the cells grew, the keratinocytes gradually metastasized while the melanocytes continued to divide, and the cell culture became more pure. (b) Identification of isolated rabbit melanocytes by L-DOPA staining. The 3^rd^ generation melanocytes were treated with L-DOPA to detect the distribution of brown or black particles in isolated cells. (c) Real-time PCR was used to detect the expression of melanocyte-specific genes such as MITF, TYR, and TYRP1 in isolated cells. (d) Isolated rabbit melanocytes were identified by immunocytochemical staining (100×) using melanocyte-specific marker proteins S-100, TYR, and TYRP1 to analyze the expression pattern of these three proteins in the isolated cells.

### Analysis of Slc7a11 gene expression in Rex Rabbit skin with different fur colors

The Slc7a11 cDNA sequence, including 31 bp 5’ UTR, 1509 bp open reading frame (ORF), and 132 bp 3’ UTRs (poly-[A] tail included), was obtained using RACE and cloning techniques and submitted to GenBank (Accession number KY971639.1) (Figure 2a). The localization of the Slc7a11 protein in the skin tissue of Rex Rabbits was determined by immunohistochemistry. Blue positive reactions were detected in the epidermis, hair bulbs, and hair root-sheaths, with different shades of color, suggesting Slc7a11 was widely expressed (Figure 2b). It was found that the expression level of the Slc7a11 gene was highest in skin with PY color, which was 3.7 times that seen in WH. The differences between PY and WH, as well as between PC and WH, were significant (*P*<0.01) (Figure 2c). Wes system analysis showed that the Slc7a11 protein was expressed in all skin tissues. The protein expression level was the highest in PY skin, and the lowest in the WH (Figure 2d). The expression levels of mRNA and protein of Slc7a11 in Rex Rabbit skin with different fur colors showed a significant positive correlation (R=0.874, *P*<0.05).

**Fig. 2.**
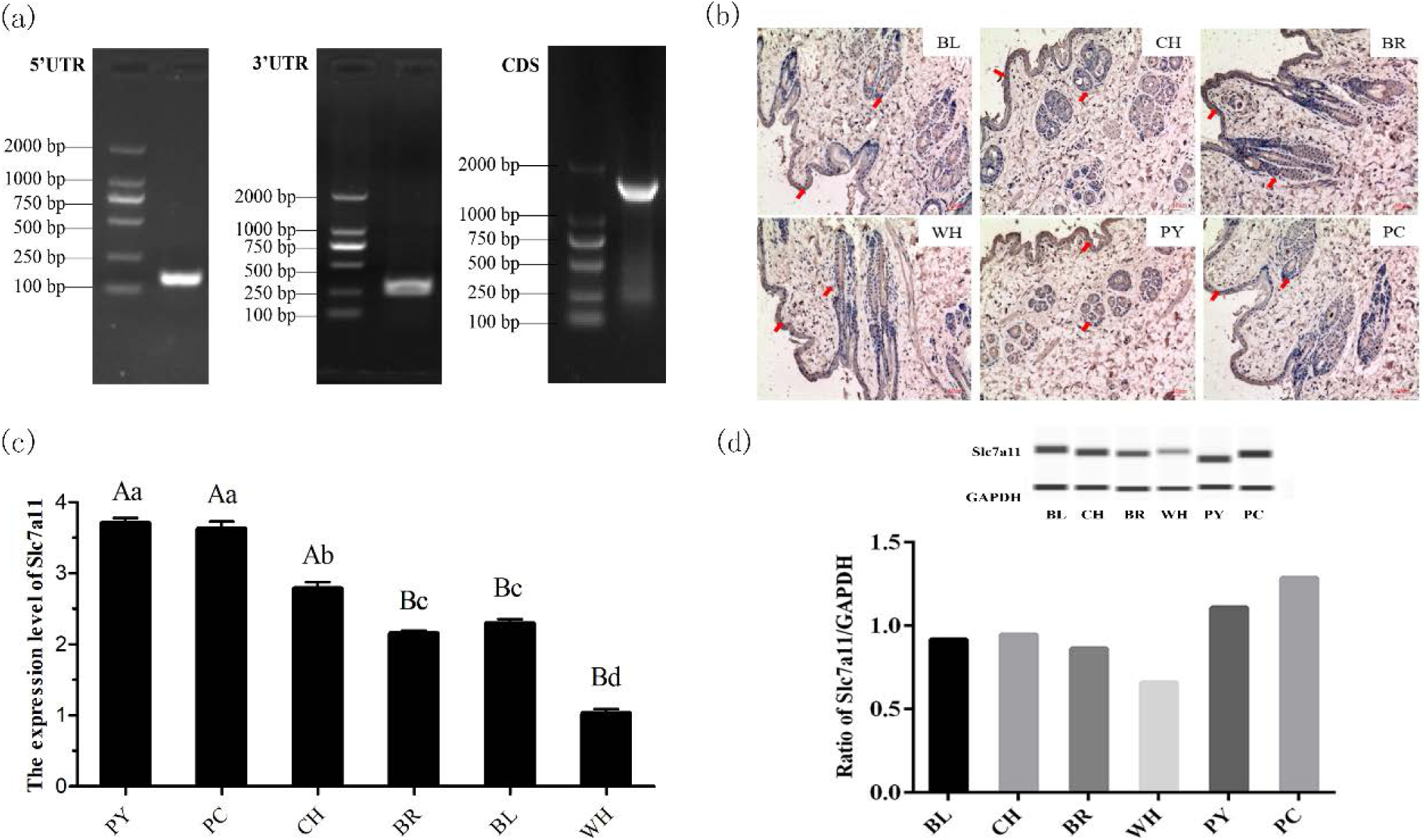
Cloning of rabbit *Slc7a11* gene and its expression in Rex Rabbit coat with different fur colors. (a) Full length sequence of rabbit *Slc7a11* cDNA was obtained using the RACE technique. The 5’UTR and 3’UTR sequences were obtained by 5’ RACE and 3’ RACE, respectively, and the DNASTAR program was used to assemble the sequence as well as remove redundant sequences to obtain the Rex Rabbit *Slc7a11* cDNA sequence. (b) Localization of *Slc7a11* in the skin of Rex Rabbits with different fur color using immunohistochemical staining. Arrows indicate positive expression of *Slc7a11* in the epidermis and hair follicles (×100). (c) mRNA expression level of *Slc7a11* gene in skin tissues of Rex Rabbits with different fur colors by Real-time PCR. (d) The expression level of *Slc7a11* (xCT) in skin tissues of Rex Rabbits with different fur colors by the Wes method. The results were analyzed using the Compass program and the relative expression ratio of *Slc7a11* was calculated.

### Effect of Slc7a11 gene expression on melanin deposition

In order to further analyze the mechanism of the Slc7a11 gene in melanogenesis, Slc7a11 siRNA interference and overexpression were performed in melanocytes, and RT-PCR and Wes were used to detect mRNA and protein expression levels of pigment-related genes such as MITF and TYR. The results showed that siRNA-2 and siRNA-3 interferences were significantly lower than that of the blank group (*P*<0.05), and siRNA-3 had the best effect (Figure 3a, 3b). pEGFP-N1-Slc7a11 was expressed in melanocytes and the expression of Slc7a11 was significantly increased (*P*<0.01) (Figure 4a, 4b).

**Fig. 3.**
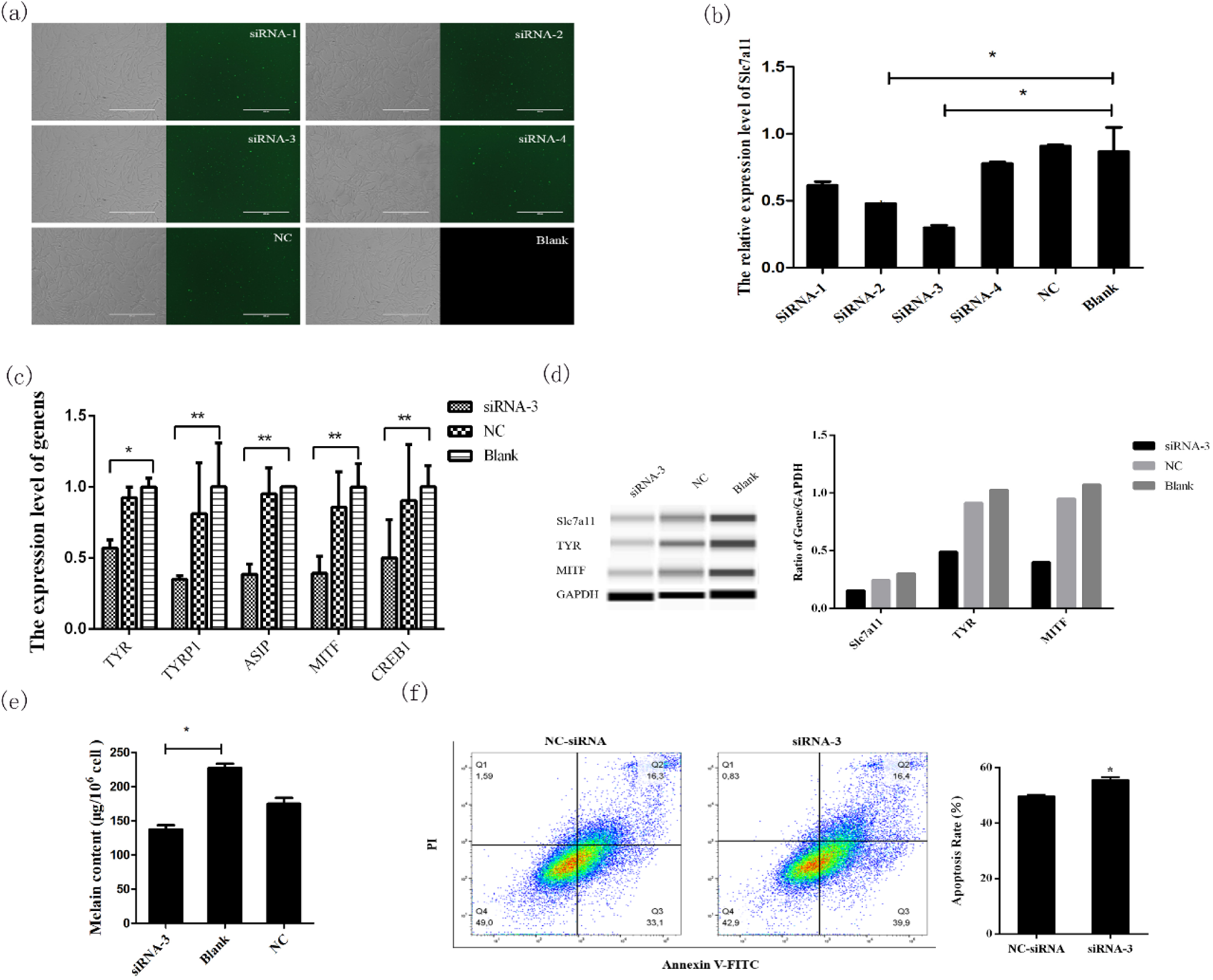
Melanogenesis-related gene expression and melanogenesis were inhibited by Slc7a11 knockdown. (a) Cell morphology 24 hours after transfection of melanocytes by FAM-siRNA. Melanocytes in the logarithmic growth phase were transfected and the transfection was detected by the observation of green fluorescence. (b) Real-time PCR detection of *Slc7a11* mRNA expression after siRNA interference. The best siRNA was screened for subsequent experiments. (c) Effects of *Slc7a11* interference on the expression of pigmentation-related genes such as MITF, TYR, TYRP1, CREB1, and ASIP. (d) Detection of the expressions of MITF, TYR, and Slc7a11 (xCT) proteins in melanocytes by Wes. The relative expression levels of MITF, TYR, and Slc7a11 (xCT) proteins were calculated and analyzed by the Compass program. (e) The effect of *Slc7a11* knockdown on melanogenesis in melanocytes. Melanocytes were collected after siRNA-3 transfection and the melanin level was measured using a microplate reader. (f) The melanocytes apoptosis rate was determined after the knockdown of Slc7a11.

**Fig. 4.**
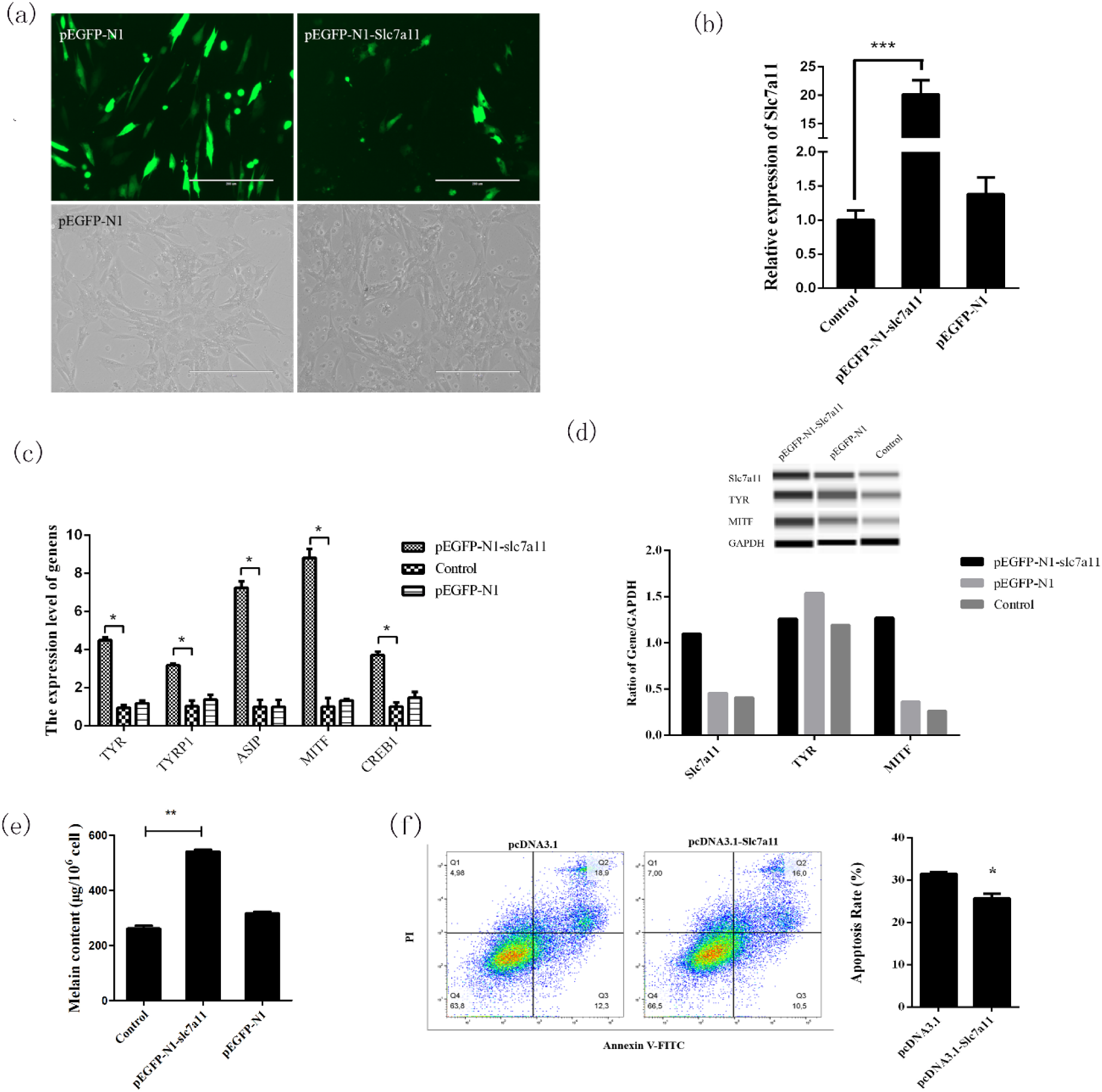
The expression of melanogenesis-related genes and melanogenesis were increased by the overexpression of Slc7a11. (a) Fluorescence detection results of pEGFP-N1-Slc7a11 transfected melanocytes. pEGFP-N1 was used as a control. (b) Detection of the overexpression of *Slc7a11* in melanocytes by Real-time PCR. (c) The effect of *Slc7a11* overexpression on the expression of pigmentation-related genes such as MITF, TYR, TYRP1, CREB1, and ASIP. (d) Detection of the expressions of MITF, TYR, and Slc7a11 (xCT) proteins in melanocytes by Wes. The relative expression levels of MITF, TYR, and Slc7a11 (xCT) proteins were calculated and analyzed using the Compass program. (e) The effect of *Slc7a11* overexpression on melanogenesis in melanocytes. Melanocytes were collected after siRNA-3 transfection and the melanin level was measured using a microplate reader. (f) The melanocytes apoptosis rate was determined after the overexpression of Slc7a11.

When Slc7a11 was overexpressed or inhibited, the mRNA (Figure 3c, 4c) and protein expression levels (Figure 3d, 4d) of genes involved in the melanogenesis pathways (such as MITF, TYR, TYRP1, CREB1, and ASIP) also changed significantly. There was a significant positive correlation between mRNA and protein expression (*P*<0.05), which was consistent with the changes in the expression of Slc7a11. Compared with the control group, the melanin level was increased when Slc7a11 was overexpressed; whereas when Slc7a11 was inhibited melanin level decreased (Figure 3e, 4e). The results suggested that Slc7a11 affects the expression of pigmentation-related genes such as TYR and MITF, and thus affects melanogenesis by melanocytes. We examined the apoptosis rate in melanocytes after transfecting with siRNA-Slc7a11 and pcDNA3.1-Slc7a11. It was found that Slc7a11 decreased melanocytes apoptosis, but inhibition of Slc7a11 enhanced apoptosis (Figure 3f, 4f).

### Identification of the core region of the Slc7a11 promoter and key transcription factor POU2F1

In order to further reveal the regulatory mechanism of the Slc7a11 gene, the promoter sequence 2499 bp before the start codon of Slc7a11 was cloned. Firstly, by predictive analysis of potential transcription factors in the Slc7a11 promoter region, four deletion vectors (P1∼P4) were constructed. Dual-luciferase assays showed that the activities of P2 and P3 were comparable (*P*>0.05). P2 activity was significantly lower than that of P1, and the activity of P4 was significantly lower than that of P3 (*P*<0.05), indicating that the deletion of -969 ∼ -469 bp and -2469 ∼ -1969 bp decreased the activity significantly. The results suggested that there were two active regions in the Slc7a11 promoter, -969 ∼ -469 bp and -2469 ∼ -1969 bp, respectively (Figure 5a).

**Fig. 5.**
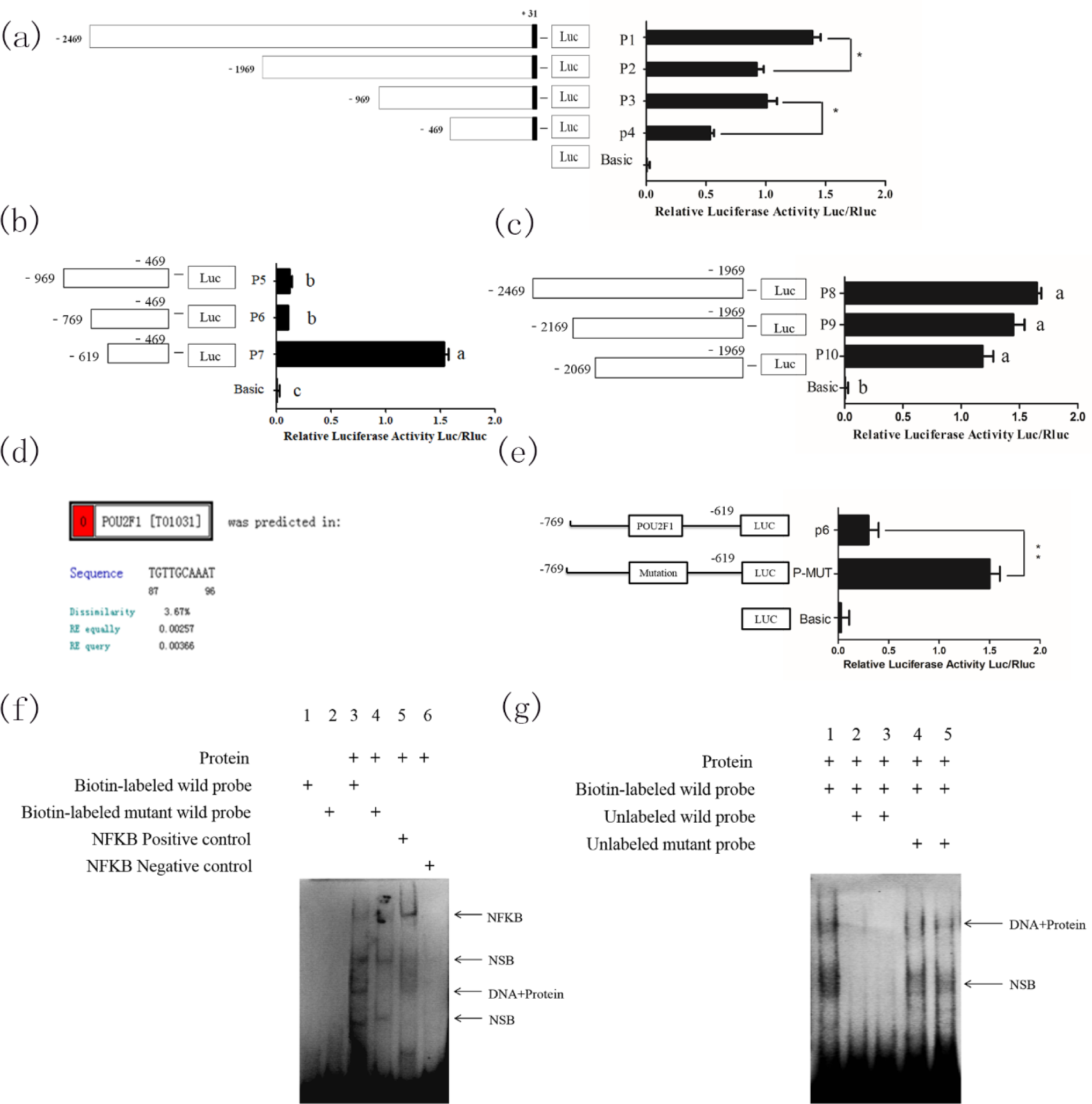
Regulation of transcriptional factor POU2F1 on *Slc7a11* promoter activity. (a) Preliminary analysis of the activity of Slc7a11 promoter-deleted vector series. P1∼P4 were constructed and the activity of each fragment was detected using dual luciferase. It was presumed that the Slc7a11 promoter contained two active regions, namely -969∼-469 bp and -2469∼ -1969 bp. (b) Activity detection of a series of vectors with deletions in the -969∼-469 bp active region of Slc7a11 promoter. P5∼P7 were designed and constructed for this region. (c) Activity detection of a series of vectors with deletions in the -2469∼-1969 bp active region of Slc7a11 promoter. P8∼P10 were designed and constructed for this region. (d) Prediction of the transcriptional binding site in the Slc7a11 promoter region. The -769 to -619bp region was the primary target based on the results of (a), (b), and (c). The results also suggested that transcriptional repressors may be present in this region. The potential transcription factor binding sites were analyzed using the online program PROMO. (e) Site-directed mutagenesis analysis of POU2F1. Based on the predicted position given by PROMO, the POU2F1 binding site was effectively mutated by site-directed mutagenesis and detected by dual-luciferase assay. (f) EMSA suggested that POU2F1 could bind to the Slc7a11 core promoter region. The 1st and 2nd lanes were normal and blank mutant probes, respectively, and no bands indicated good probes. The 3rd and 4th lanes were biotin-labeled normal and mutation probes, respectively. The 5th and 6th lanes were NF-KB positive and negative controls, respectively. NSB stands for non-specific binding. (g) Specific binding of POU2F1 to the Slc7a11 core promoter region by competitive EMSA experiments. In 2nd and 4th lanes, unlabeled probes were 40 times that of the labeled probes, and in 3rd and 5th, unlabeled probes were 80 times that of the labeled probes.

In order to further identify the core transcription factor binding region, a series of deletion vectors were constructed targeting the -2469 ∼ -1969 bp and -969 ∼ -469 bp fragments: P8, P9, P10 for the -2469 ∼ -1969 bp region, and P5, P6, and P7 for the -969 ∼ -469 bp region. The activities of P8, P9, and P10 were similar by luciferase assay (*P*>0.05) (Figure 5b). However, the activities of P5 and P6 were comparable but significantly lower than that of P7, indicating that the activity was significantly reduced with the deletion of the -769 to -619 bp region (*P*<0.05) (Figure 5c), suggesting that -769 ∼ -619 bp is the core transcription region of Slc7a11. The predicted POU2F1 binding site (−713 to -703 bp) was found in the -769 to -619 bp region (Figure 5d). It was found that the promoter activity of Slc7a11 was significantly increased after the site-directed mutation (*P*<0.01), indicating that POU2F1 inhibited the promoter activity of Slc7a11 (Figure 5e).

To further determine whether POU2F1 binds to this site of the Slc7a11 promoter, an EMSA experiment was performed using the nuclear protein of melanocytes (Figure 5f). The 3rd lane showed that the biotin-labeled probe of POU2F1 could bind to the nuclear protein to form a complex band. No band in the 4th lane suggested that mutated POU2F1 could not bind to a nuclear protein to form a complex. The results together revealed that POU2F1 could bind to the Slc7a11 core promoter region. And a competitive EMSA experiment was performed to further determine whether the binding was specific (Figure 5g). A complex band in Lane 1 indicated that the probe was able to bind to the nuclear protein. The 2nd and 3rd lanes were cold-competitive reactions with unlabeled normal probes. No bands were observed, indicating that the unlabeled normal probes were competitively bound to the nuclear protein due to their large amounts, meaning the biotin-labeled probe could hardly bind to the protein. The 4th and 5th lanes were cold-competitive groups with unlabeled mutant probes. The unlabeled normal probe did not bind to the protein after mutation, and thus did not compete with the biotin-labeled normal probe, producing protein-probe complex bands. The results confirmed that the POU2F1 protein could bind to the -713 to -703 bp region and inhibit the activity of the Slc7a11 promoter.

## Discussion

Fur color is controlled by different genes in the process of pigment biosynthesis. The differences in color are mainly due to the different ratios between pheomelanin (red and yellow) and eumelanin (black) (Barsh and Cotsarelis, 2007; Ito and Wakamatsu, 2008; Ito et al., 2000). The main model currently proposed is that the ratio of eumelanin/pheomelanin in mammalian pigments is solely or indirectly regulated by the activity of tyrosinase – the rate-limiting enzyme of melanin synthesis (Wagner et al., 2001). This model inferred that in the presence of low concentrations of tyrosinase, dopaquinone reacts with cysteine to produce cysteine dopa, thereby increasing the level of pheomelanin (Ito, 2003). However, this pattern is not yet fully understood. Studies have confirmed that the xCT transporter encoded by the Slc7a11 gene is crucial for the regulation of pigments and can directly affect the increase of pheomelanin (Chintala et al., 2005). Based on previous studies, Rex Rabbits with a variety of natural fur colors were used to explore the expression pattern of Slc7a11 gene in dorsal skin tissues with different fur colors. xCT was expressed in the epidermis, hair bulb, and hair root-sheaths of the skin tissues examined by immunohistochemistry. It is known that melanocytes in the skin are often distributed in different regions when matured and that the only place where melanin can be supplied to the hair shaft is the hair bulb (Slominski et al., 2005; Tobin, 2011; Tobin et al., 1999). In this study, it was found that the expression sites of the Slc7a11 gene were consistent with the distribution of melanin, suggesting that the protein was related to the formation of Rex Rabbit fur color. Moreover, the Slc7a11 gene had the highest expression level in protein yellow coat, and the lowest level in white coat, by real-time quantitative and Wes system analyses. It is speculated that Slc7a11 affects the formation of cystine, which is reduced to cysteine, and thus alters the production of pheomelanin. This is consistent with studies on alpaca and sheep (He et al., 2012; Tian et al., 2015). Knockdown and overexpression analyses of Slc7a11 confirmed that this gene can affect the expression of TYR, MITF, TYRP1, and ASIP in the melanogenesis pathway (Gutierrez-Gil et al., 2007; Hida et al., 2009). Moreover, Slc7a11 could decrease melanocytes apoptosis and further affect the melanogenesis of melanocytes. These results confirmed that Slc7a11 is closely related to the formation of Rex Rabbit fur color. And the regulatory factors of such expression patterns would be the next research objective.

Chintala et al. found that the mouse light gray (sut) mutation was due to the inhibition of phaeomelanin production. It was caused by a large deletion of the Slc7a11 gene starting from the 11th intron crossing the 12th exon into the region adjacent to the Pcdh18 gene. This resulted in a change in the 3’ end of Slc7a11 transcription(Chintala et al., 2005). Based on these results, the identification of similar deletions in the natural populations of Rex Rabbits with six fur colors was carried out. Unfortunately, no such large fragment deletions were found in similar areas. Exon scanning did not show any SNPs sites, indicating that Slc7a11 is relatively conserved in the population. Similar results have been seen in humans and sheep (Gasol et al., 2004; He et al., 2012; Sato et al., 2000). The regulatory mechanism of Slc7a11 is unkonwn.

Further, the promoter series deletion vector dual-luciferase was used to search for the -769 to -619 bp transcription core region of Slc7a11. And it’s confirmed that POU2F1 protein binds to the -713 to -703 bp region of the Slc7a11 promoter to inhibit its activity. POU2F1, also known as Octamer Transcription factor-1 (Oct-1), is a widely expressed POU protein factor. Recent studies suggest that it can regulate target genes associated with processes such as oxidation, anti-cytotoxicity, stem cell function, and cancer development, etc. (Maddox et al., 2012; Vazquez-Arreguin and Tantin, 2016). Ethanol has been reported to increase the expression of Slc7a11 by reducing the binding of POU2F1 to the Slc7a11 gene promoter (Lin et al., 2013). In this study, POU2F1 was found to specifically bind to the Slc7a11 promoter and inhibit its transcription. Together with the previous finding that Slc7a11 promotes melanin cytochrome deposition, POU2F1 can be used as a target for artificial modification of animal fur colors.

## Conclusions

In summary, rabbit melanocytes were isolated and identified. The Slc7a11 expression levels in the protein yellow colored skin tissue was higher than those with other fur colors. It was verified that Slc7a11 could significantly affect the protein and mRNA expressions of TYR and MITF, inhibiting melanocytes apoptosis, thus affecting the melanogenesis by melanocytes. Further, it’s confirmed that POU2F1 regulated the activity of the rabbit Slc7a11 promoter. Our results provided a theoretical basis for further exploration of the role Slc7a11 plays in pigmentation.

## Abbreviations

Slc7a11: (Solute carrier family 7 member 11);
POU2F1: (POU domain class 2 transcription factor 1);
RNA-Seq: (transcriptome sequencing);
BL: (black);
CH: (chinchilla);
WH: (white);
BR: (brown);
PY: (protein yellow);
PC: (protein chinchilla);
EMSA: (Electrophoresis mobility shift assay);
SD: (standard deviation);
ORF: (open reading frame);
Oct-1: (Octamer Transcription factor-1).

## Acknowledgements

This work was supported by the Modern Agricultural Industrial System Special Funding (CARS-43-A-1), National Natural Science Foundation of China (Grant No. 31702081), and Science and Technology Major Project of New Variety Breeding (Livestock and Poultry) of Zhejiang Province, China (2016C02054-10).

## Statement of Ethics

This study was carried out in accordance with the recommendations of Animal Care and Use Committee at Yangzhou University. The protocol was approved by the Animal Care and Use Committee at Yangzhou University.

## Author Contributions

Yang Chen conceived and designed the experiments, performed the experiments, wrote the paper. Shuaishuai Hu performed the experiments. Lin Mu prepared figures and/or tables. Bohao Zhao analyzed the data. Manman Wang and Naisu Yang contributed reagents/materials/analysis tools. Guolian Bao and Cigen Zhu prepared figures and/or tables. Xinsheng Wu conceived and designed the experiments, reviewed drafts of the paper.

